# Evaluation of Precision of the *Plasmodium knowlesi* Growth Inhibition Assay for *Plasmodium vivax* Duffy-Binding Protein-based Malaria Vaccine Development

**DOI:** 10.1101/2024.01.23.576905

**Authors:** Jonas E. Mertens, Cassandra A. Rigby, Martino Bardelli, Doris Quinkert, Mimi M. Hou, Ababacar Diouf, Sarah E. Silk, Chetan E. Chitnis, Angela M. Minassian, Robert W. Moon, Carole A. Long, Simon J. Draper, Kazutoyo Miura

## Abstract

Recent data indicate increasing disease burden and importance of *Plasmodium vivax* (*Pv*) malaria. A robust assay will be essential for blood-stage *Pv* vaccine development. Results of the *in vitro* growth inhibition assay (GIA) with transgenic *P. knowlesi* (*Pk*) parasites expressing the *Pv* Duffy-binding protein region II (*Pv*DBPII) correlate with *in vivo* protection in the first *Pv*DBPII controlled human malaria infection (CHMI) trials, making the *Pk*GIA an ideal selection tool once the precision of the assay is defined. To determine the precision in percentage of inhibition in GIA (%GIA) and in GIA_50_ (antibody concentration that gave 50 %GIA), ten GIAs with transgenic *Pk* parasites were conducted evaluating four different anti-*Pv*DBPII human monoclonal antibodies (mAbs) at different concentrations, and three GIAs were conducted testing eighty anti-*Pv*DBPII human polyclonal antibodies (pAbs) at 10 mg/mL. A significant assay-to-assay variation was observed, and the analysis revealed a standard deviation (SD) of 13.1 in the mAb and 5.94 in the pAb dataset for %GIA, with a LogGIA_50_ SD of 0.299 (for mAbs). Moreover, the ninety-five percent confidence interval (95%CI) for %GIA or GIA_50_ in repeat assays was calculated in this investigation. These results will support the development of future blood-stage malaria vaccines, specifically second generation *Pv*DBPII-based formulations.

## 1. Introduction

Malaria is the most pernicious of parasitic diseases and exerts an enormous health and socioeconomic burden on many of the most vulnerable populations and poorest regions on earth, in recent years only aggravated by the impending climate crisis [1]. Six protozoan parasite species of the genus *Plasmodium* are known to infect humans, among them, *Plasmodium falciparum* (*Pf*) and the lesser studied *Plasmodium vivax* (*Pv*) are responsible for the bulk of these infections [1]. While the lion’s share of morbidity and mortality is caused by *P. falciparum* and is mostly confined to Sub-Saharan Africa, *P. vivax* is found more extensively, with *Pv* infections occurring in most tropical and subtropical regions of the world, thus responsible for the majority of cases outside Sub-Saharan Africa [2].

While the overall lifecycles of the two species are similar, key biological differences impact epidemiology and complicate *P. vivax* malaria control [3,4], and can be considered as being factors in this parasite’s different and more widespread distribution. Firstly, the earlier production of *Pv* gametocytes leads to a more rapid transmission. Secondly, *Pv* possesses the ability to form dormant parasites within the liver, called hypnozoites, which can reactivate weeks, months or even years after primary infection, facilitating waves of relapsing parasitaemia, illness, and transmission [5]. Remarkably, these relapses have been estimated to account for up to 80 – 90 % of new infections [6]. Thirdly, *Pv* merozoites exhibit a tropism for Duffy antigen receptor of chemokines (DARC, also known as Fy glycoprotein (Fy)) positive red blood cells (RBCs), thus limiting endemicity in West and Central African populations, where Duffy blood group negativity provides natural resistance against *Pv* infection [7]. Moreover, *Pv* shows a clear tropism for reticulocytes, for which reticulocyte binding-like proteins (RBPs) appear to be responsible [8]. Consequently, 3.3 billion humans were living under the risk of *Pv* transmission in 2017 [9], especially in the Americas, Oceania (particularly the Western Pacific), South East Asia, and the Eastern Mediterranean [1]. In most of these regions, *P. vivax* is the most prevalent malaria parasite [1] with recent data demonstrating a significant burden of morbidity and associated mortality in young children and pregnant women [10]. Termed “benign” for many years, this resilient parasite species in fact carries all of the attributes of “perniciousness” historically only linked to *Pf* [11]. Additionally, in regions where *Pv* and *Pf* are co-endemic and *Pf* infection risk has been lessened by control measures, there is a converse risk increase for *Pv* infection; this increase is also seen in patients treated for *Pf* malaria in these areas [4]. Therefore, ignorance of *Pv* seems irrational.

Historically, efforts to develop a *Pv* vaccine have lagged behind *Pf* because of critical bottlenecks in the development process [12], among them the absence of well-characterized anti-parasitic functional assays due to the lack of a long-term *in vitro* culture system. Hence only few novel candidate vaccines are in the pipeline or have even progressed to the clinic [12]. Nevertheless, recent breakthroughs in *Pv* vaccine development have been realized. Building on prior work in Australia [13], a safe *Pv* blood-stage controlled human malaria infection (CHMI) model was established for the first time in Europe with a Thai parasite clone termed *Pv*W1, whose genome was reported with a high quality assembly [14]. Moreover, two vaccines targeting the leading *P. vivax* blood-stage antigen *Pv* Duffy-binding protein region II (*Pv*DBPII), based on *Pv* strain Salvador-1 (Sal-1) sequence, have advanced along the clinical development pipeline [15,16]. The *Pv*DBPII contains the receptor binding domain which interacts with the DARC found on reticulocytes [17], thus facilitating parasite invasion of these RBCs [18].

A major milestone was next met when the protein *Pv*DBPII vaccine formulated in Matrix-M™ adjuvant showed ∼50 % reduction in *Pv* parasite growth in the blood of vaccinees following CHMI with the *Pv*W1 clone of *P. vivax* [19]. This *Pv*WI clone used for CHMI harbours a single copy of the *Pv*DBP gene with a heterologous sequence to the recombinant Sal-1 *Pv*DBPII protein employed for vaccination, i.e. protection was afforded in heterologous challenge [19]. In addition, the A1-H.1 strain of the zoonotic *Plasmodium knowlesi* (*Pk)* species has been adapted for long-term *in vitro* culture in human RBCs and most importantly, transgenic *Pk* parasites expressing *Pv*DBP have been developed [20– 22]. As *Pv*DBP is able to fully complement the essential role of its *Pk* orthologue in erythrocyte invasion, these parasites thus provide the first means to routinely screen for functional anti-parasitic antibody activity *in vitro*, without necessitating access to *Pv*-infected blood from the field [23]. These *Pk*-*Pv*DBP parasites have now enabled functional screening of anti-*Pv*DBPII mAbs [20,24], whilst human purified total IgG from *Pv*DBPII vaccinees in the CHMI clinical trial showed functional *in vitro* growth inhibition that correlated with the *in vivo* growth inhibition () of *Pv*W1 parasites [19].

Future work will seek to build on these recent findings. Notably, for the development of a next-generation vaccine, a tool for facilitation of preclinical and clinical go/no-Go decisions with regards to vaccine candidate selection will be essential. *In vitro* growth inhibition assays (GIAs) for assessment of antibody-driven effects on parasite invasion or growth have been an integral part of blood-stage *P. falciparum* malaria research for many years. Notably, GIAs that asses inhibitory activity of antibodies using transgenic *Pk* parasites expressing *Pv*DBP could now fill the exigent role of a candidate selection tool for improved *Pv* vaccines; thereby defining novel immunogen designs and/or formulations that elicit significantly higher levels of growth inhibition in GIAs than the benchmark *Pv*DBPII/Matrix-M™ vaccine. In turn, these formulations would be predicted to facilitate much greater levels of IVGI in humans and ultimately, full protection. For this, a robust assay, which can provide reliable and biologically relevant data with high precision, is of paramount importance. Generally speaking, GIAs differ with regards to the parasite species and strain as well as methodology employed, therefore each iteration of assay must be evaluated individually. With another investigation having just tested the precision or “error of assay” (EoA) in *Pf*GIA readouts for a reticulocyte-binding protein homologue 5 (RH5)-based *Pf* vaccine by assessing parasite lactate dehydrogenase activity (pLDH) [25], our study reported here now characterizes the EoA in the aforementioned *Pk*GIA using monoclonal and polyclonal antibodies against the *Pv*DBPII.

## 2. Materials and Methods

### 2.1 Plasmodium knowlesi (Pk) parasite culture and synchronization at the University of Oxford

Development of transgenic *Pv*DBPOR/Δβγ parasites was previously reported. In brief, these represent *P. knowlesi* parasites of the parental A1-H.1 strain which were genetically modified to express *P. vivax* Salvador-1 (Sal-1) strain *Pv*DBP in place of the native *Pk*DBPα. Using CRISPR-Cas9 genome editing, the *Pk*DBPα gene was replaced by the *Pv*DBP orthologue (OR) with subsequent deletion of the *Pk*DBPβ and *Pk*DBPγ paralogues, thereby creating a transgenic *P. knowlesi* line reliant on the *Pv*DBP for invasion of erythrocytes [21]. Parasites were cultured in type O, Rh+ blood from different human donors, obtained both in-house from volunteers at the University of Oxford and from the United Kingdom’s National Health Service Blood and Transplant (NHSBT). Fy serophenotyping was done using anti-Fy(a) monoclonal, anti-Fy(b) polyclonal andanti-human IgG/anti-human globulin blood typing reagents (Lorne Laboratories). The cultures were set up and maintained according to previously described protocols [23]. For maintenance, cultures were incubated at 37 °C in non-vented flasks containing an atmosphere with a gas mixture of 5 % O_2_, 5 % CO_2_ and 90 % N_2_. The incomplete *Pk* culture medium was prepared using 500 mL of RPMI-1640 liquid medium (Sigma-Aldrich R0883) to which 2.97 g HEPES (Sigma-Aldrich H3375), 0.025 g hypoxanthine (Sigma-Aldrich H9636), 0.15 g NaHCO_3_ (Sigma-Aldrich S5761), 1 g D-glucose (Sigma-Aldrich G7021) and 10 mL 100X L-glutamine (Gibco 25030) were added. To complete the medium, 10 mL pooled heat-inactivated filter-sterilized human O+ serum obtained from NHSBT was mixed with 40 mL *Pk* incomplete culture medium and 50 μL 10 mg/mL gentamicin (Sigma-Aldrich G1272). The 2x *Pk* complete medium used in the GIAs was prepared by mixing 30 mL *Pk* incomplete culture medium with 20 mL pooled heat-inactivated filter-sterilized human O+ serum and 100 μL 10 mg/mL gentamicin. If sufficient late-stage parasites (i.e. > 2 % parasitaemia) were present in a culture, synchronisation at trophozoite or schizont stage was performed by utilizing magnetic activated cell sorting (Miltenyi Biotec MACS LD columns).

### 2.2 Monoclonal antibody (mAb) production and purification

The anti-*Pv*DBPII human IgG1 mAbs DB1, DB5, DB6 and DB9 [20] were produced by transient transfection of HEK Expi293 cells (Thermo Fisher Scientific). Briefly, cells were transfected following the manufacturer’s protocol using ExpiFectamine™ (Thermo Fisher Scientific), including the addition of enhancer 1 and enhancer 2 (Thermo Fisher Scientific) 18 h post-transfection. Supernatants were harvested seven days after transfection via centrifugation and mAbs were purified from culture supernatants using a 5 mL protein G column (Cytiva) in Tris-Buffered Saline (TBS) on a fast protein liquid chromatography (FPLC) system (Cytiva ÄKTA Pure). The mAbs were eluted as 1.7 mL fractions in glycine (200 mM, pH 2.4) then neutralized with Tris buffer (1 M, pH 9.0). These fractions were then pooled and concentrated to 10 mL before size exclusion chromatography (SEC) purification using a SEC column (Cytiva Superdex 200 HiLoad 16/600 column) on the FPLC system into TBS. Finally, mAbs were concentrated and buffer-exchanged into incomplete *Pk* medium for the use in GIAs.

### 2.3. Growth inhibition assays (GIAs)

#### 2.3.1 GIAs with *Pk* parasites at the University of Oxford

Measurement of growth inhibition activity was adapted for *Pk* from protocols from the Laboratory for Malaria and Vector Research (LMVR) at National Institute of Allergy and Infectious Disease (NIAID), National Institutes of Health (NIH), United States of America [26]. After dilution to the desired concentrations with incomplete *Pk* medium, 20 μL of the mAb samples and controls were introduced into sterile 96-well flat/half area tissue culture plates (Corning 3696) in triplicates. The mAbs were tested at concentrations of 2, 0.4, 0.08 and 0.016 mg/mL (DB1, DB5 and DB6) and 2, 0.8, 0.4, 0.125, 0.08, 0.04 and 0.016 mg/mL (DB9), respectively. When synchronization was complete, trophozoite cultures were diluted to a late-stage parasitaemia of 1.5 % at 4 % haematocrit in 2x *Pk* complete medium and then pipetted in volumes of 20 μL into aforementioned 96-well plates. Control wells included: only infected erythrocytes and culture medium (normal parasite growth); infected erythrocytes incubated in the presence of 5 mM EDTA (total inhibition of parasite growth); and infected erythrocytes plus the anti-Ebola virus glycoprotein human IgG1 antibody EBL040 [27] (negative control mAb, no inhibition of parasite growth). The plates were incubated at 37 °C for ∼27 h, equivalent to one lifecycle of *Pk* in vitro. Afterwards, parasite growth in every well was evaluated using pLDH activity. For use in the assay, 500 mL LDH buffer solution consisting of 50 mL 1M Tris HCl (pH 8.0, Sigma-Aldrich T3038) and 450 mL ddH_2_O, to which 2.8 g sodium L-lactate (Sigma-Aldrich L7022), and 1.25 mL Triton X-100 (Sigma Aldrich X100) were added and mixed for at least 30 min, were prepared. Subsequently, a 10 mg nitro blue tetrazolium (NBT) tablet (Sigma-Aldrich N5514) was introduced to 50 mL of this mixture. Just prior to assay development, 50 μL 10 mg/mL 3-acetylpyridine adenine dinucleotide (APAD; Sigma-Aldrich A5251) and 200 μL 50 U/mL diaphorase (Sigma-Aldrich D5540) were added to every 10 mL LDH buffer/nitro blue tetrazolium mixture. 120 μL of this mixture was then added to every well. Plates were read with a microplate reader (BioTek TS800 absorbance reader) and the accompanying software (BioTek Gen5 software) at 650 nm once the optical density had reached 0.4 to 0.6 in the infected erythrocyte/medium control wells. Percentage of growth inhibition in the growth inhibition assay (%GIA) was then calculated using the following formula:

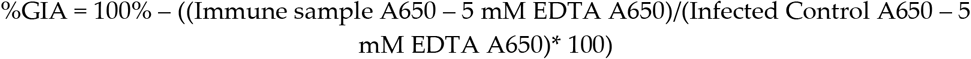

#### 2.3.2 GIA with *Pk* parasites at the LMVR

At the LMVR, human polyclonal antibodies (pAbs), which were collected from three Phase I/IIa clinical trials called VAC069, VAC071 and VAC079, were evaluated. As reported previously [19], these trials received ethical approval from UK National Health Service Research Ethics Services, (VAC069: Hampshire A Research Ethics Committee, Ref 18/SC/0577; VAC071: Oxford A Research Ethics Committee, Ref 19/SC/0193; VAC079: Oxford A Research Ethics Committee, Ref 19/SC/0330). The vaccine trials were also approved by the UK Medicines and Healthcare products Regulatory Agency (VAC071: EudraCT 2019-000643-27; VAC079: EudraCT 2019-002872-14). All participants provided written informed consent and the trials were conducted according to the principles of the current revision of the Declaration of Helsinki 2008 and ICH guidelines for Good Clinical Practice. The methodology of the *Pk*GIA and median %GIA value from three independent *Pk*GIA for each pAb have been published elsewhere [19]. In this study, the same data were reanalysed to determine the EoA of *Pk*GIA at the LMVR. In brief, the *Pk*GIA was performed at 10 mg/mL purified total IgG (Protein G purified from serum) by mixing with ∼1.5 % trophozoite-rich parasites in a final volume of 40 μL in 96-well plates. After ∼27 h of incubation, the relative parasitaemia in each well was determined by pLDH activity.

### 2.4. Statistical analysis

For correlation analyses, a Spearman rank test was utilized (GraphPad Prism software, version 9.3.1) and *p* < 0.05 was considered as significant. The other analyses were performed using R (version 4.2.1, The R Foundation for Statistical Computing). To evaluate total variance and sources of variance (either variance that was determined by test antibody and test concentration, or residual of variance) in %GIA, linear model fits were performed using the lm function. Based on the residual variance, standard deviation (SD) of %GIA was calculated. To determine the 95 percent confidence interval (95%CI) of the SD in %GIA readout, assay-stratified bootstrap analysis was performed, where a data set was stratified by the assay number first; then resampling was performed by the assay number instead of individual data points because of the occurrence of significant assay-to-assay variation. The 95%CI of SD was estimated from 1,000 replications. For each mAb in each assay, the antibody concentrations that gave 50, 40 or 30 %GIA (GIA_50_, GIA_40_, GIA_30_, respectively) was calculated using a four-parameter logistic model with the lower asymptote parameter fixed at 0 using the L.4 function in the drc package version 3.0-1. The SD and 95%CI of SD for Log-transformed GIA_50_, GIA_40_, or GIA_30_ readouts were calculated as above.

## 3. Results

At the University of Oxford, A1-H.1 *P. knowlesi* malaria parasites expressing Salvador-1 (Sal-1) strain *Pv*DBP (*Pv*DBPOR/Δβγ) instead of their native *Pk*DBPα were cultured in human RBCs from different donors and used in ten GIAs. In these GIAs, four human IgG1 mAbs (anti-*Pv*DBPII antibodies DB1, DB5, DB6 and DB9) were tested with eight different batches of RBCs on eight different days. Testing of growth inhibition was done at concentrations of 2, 0.4, 0.08 and 0.016 mg/mL for DB1, DB5 and DB6, as well as 2, 0.8, 0.4, 0.125, 0.08, 0.04 and 0.016 mg/mL for DB9 in each assay. Additionally, as a negative control, EBL040 (an anti-Ebola virus human IgG1 mAb [27]) was used at a concentration of 0.5 mg/mL. For each anti-*Pv*DBPII mAb, at each test concentration, average (Avg), standard deviation (SD) and percentage of coefficient of variation (%CV) in percentage of inhibition in GIA (%GIA) were calculated from the ten assays (**Figure 1**). The original GIA values can be found in supplementary **Table S1**. To determine an appropriate model for EoA analysis, correlations between Avg and SD, or between Avg and %CV, were evaluated. There was no obvious effect by the different mAbs used on either SD or %CV. Similar to what was seen in an earlier publication, where *Pf*GIAs were conducted for one of the leading blood-stage antigens for *P. falciparum*, the reticulocyte-binding protein homologue 5 (RH5) [25], the SD was relatively stable with no significant correlation between Avg and SD (*p* = 0.581). Regarding %CV on the other hand, there was a strong negative correlation between Avg and %CV, i.e. %CV decreased with increasing Avg (*p* < 0.0001 by a Spearman’s rank test). Hence, the further analysis conducted was based upon a constant SD model with non-transformed %GIA.

**Figure 1:**
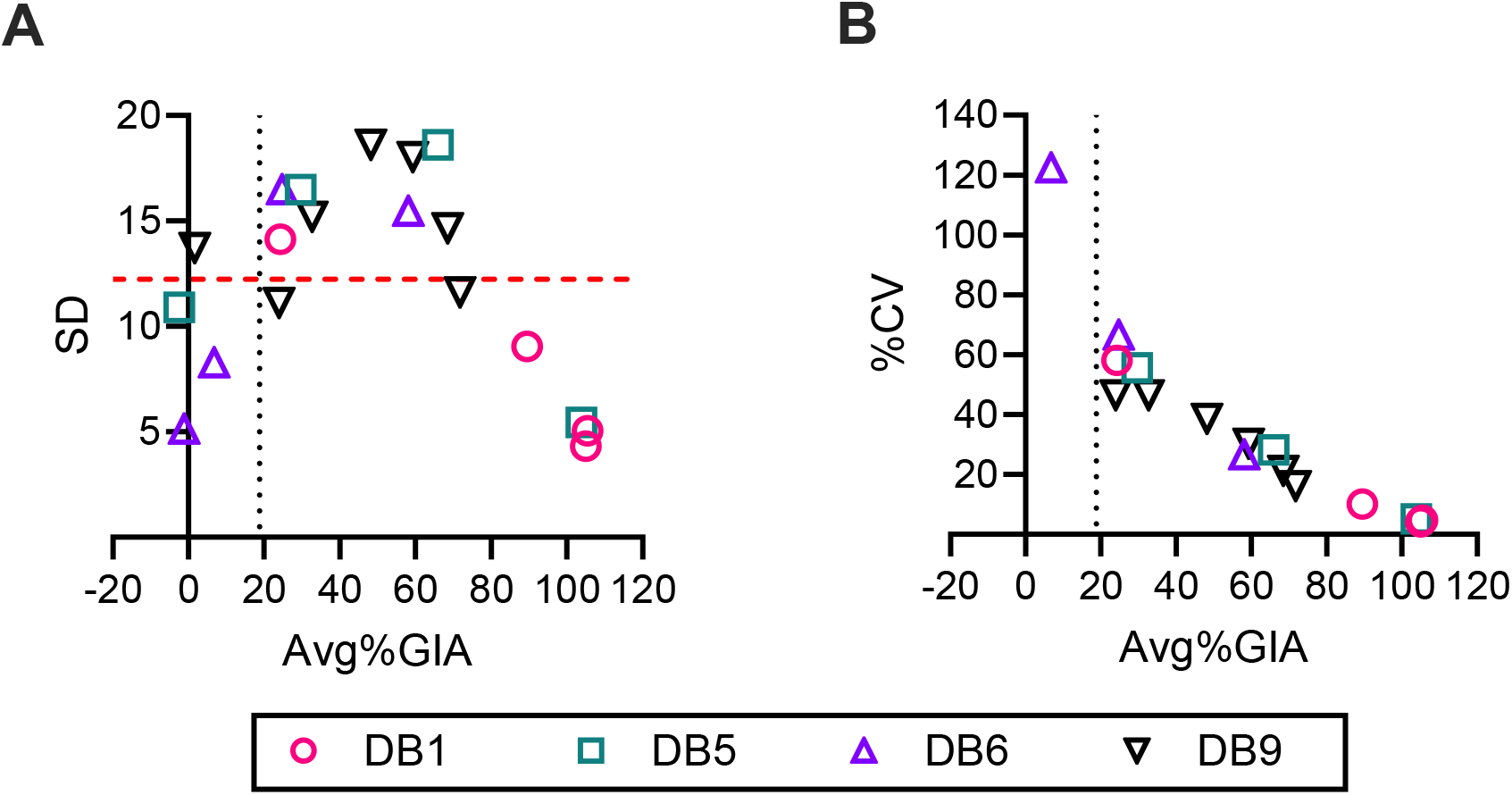
Comparison of SD vs %CV in *Pk*GIA with four different mAbs. *Plasmodium knowlesi* parasites genetically modified to express the *P. vivax* Sal-1 PvDBP (*Pv*DBP^OR^/Δβγ) were cultured and then used in GIAs to test the growth-inhibition activity of four anti-*Pv*DBPII mAbs (DB1, DB5, DB6, DB9). The antibodies were evaluated at concentrations of 2/0.4/0.08/0.016 mg/mL (DB1, DB5, DB6) and 2/0.8/0.4/0.125/0.08/0.04/0.016 mg/mL (DB9). From ten GIAs, average (Avg), standard deviation (SD) and percentage of coefficient of variation (%CV) were calculated. Results for Avg vs SD and Avg vs %CV (**B**) are shown. The vertical black line in both panels indicates the mean +2 SD value for 0.5 mg/mL EBL040 (negative control mAb). The horizontal red line in (**A**) demonstrates the mean SD (12.2) of all data points. Three data points with an Avg%GIA value between -2and 2 %GIA (absolute %CV > 400) in (**B**) are not shown.

For determination of the EoA in %GIA, the difference in measured %GIA from Avg%GIA was calculated for each mAb at each concentration in every assay (ΔAvg%GIA). A strong assay effect was observable on ΔAvg%GIA (**Figure 2**). For instance, in assays 01 (A01) or 06 (A06), the majority of data points showed negative ΔAve%GIA values (i.e., lower %GIA than the average of all ten assays). Conversely, the majority of data points in A03 or A10 were of higher %GIA than the ten-assay-average. A linear regression analysis was thus conducted, in which ΔAvg%GIA was utilized as a response variable. The specific mAb, the test concentration of the mAb and one of three factors (assay day (8 different days), assay number (A01 – A10), and RBC number (R01 – R08)) were included as explanatory values in each analysis. In all three regression analyses undertaken, the specific mAb and test concentration did not have significant impact (*p* > 0.999) on ΔAvg%GIA, indicating that EoA was similar among different mAbs at different test concentrations. On the other hand, the impact of assay day, assay number and RBC number on EoA were highly significant (*p* < 0.001). The variation due to Duffy blood group serophenotype (Fy) was difficult to evaluate in this study, because no single assay evaluated two or three Fy serophenotypes simultaneously. However, serophenotype-to-serophenotype variation in ΔAvg%GIA seems smaller than the assay-to-assay variation seen in the six assays (A01, A02, A03, A04, A06 and A07) where all assays were conducted using RBCs with the same Fy^a-/b+^ serophenotype.

**Figure 2:**
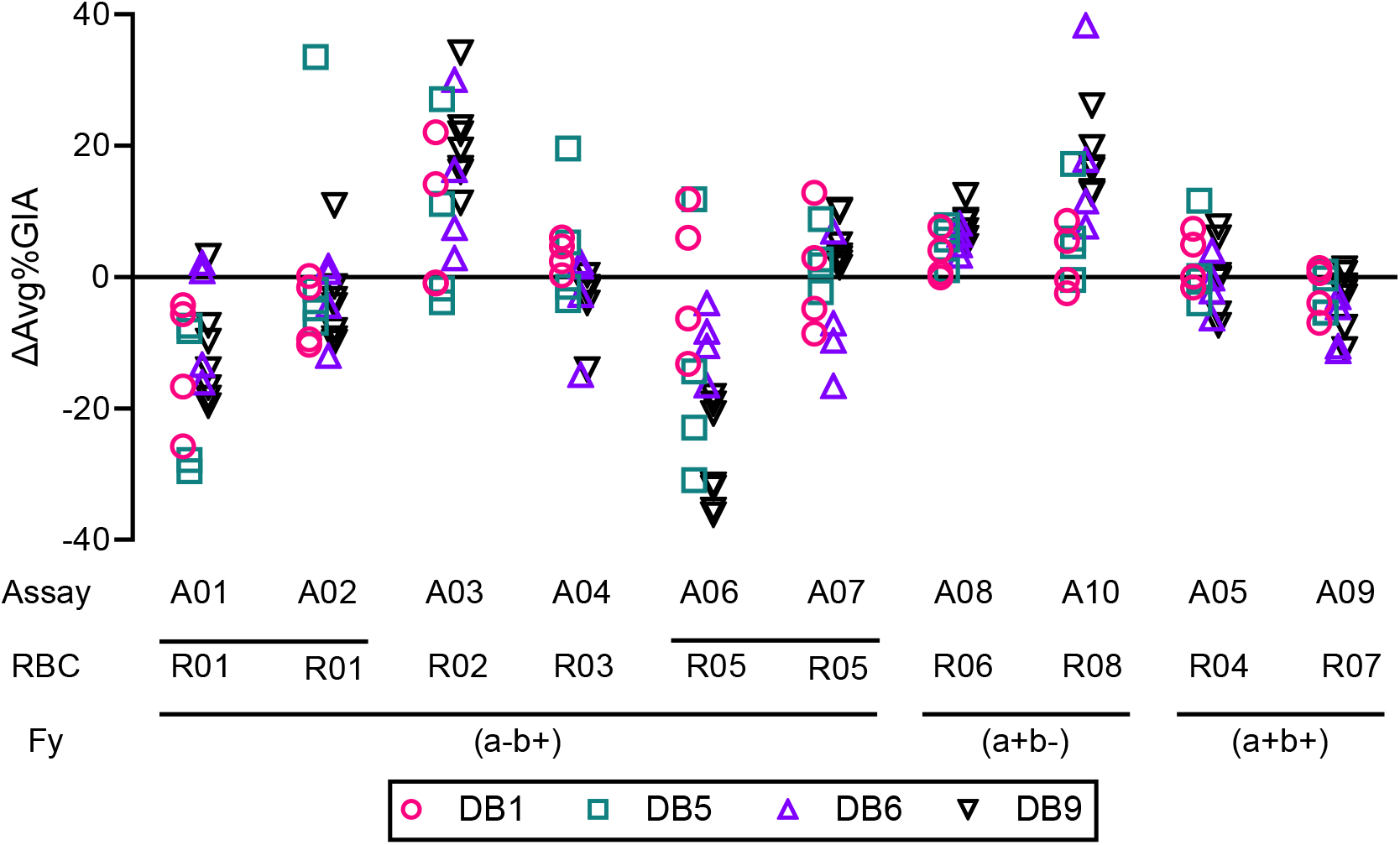
EoA in %GIA by ΔAvg%GIA for all 10 assays conducted. The mAbs were tested in 10 independent assays (A01 to A10) using 8 different batches of RBCs (R01 to R08). In each assay, the difference from Avg%GIA (ΔAvg%GIA) was determined for each mAb at each concentration. Assays A01, A02 and A03 were done on the same day, all other assays were performed on different days. The results are grouped by Duffy blood group serophenotype (Fy) of the RBCs employed in each assay.

Another linear regression analysis was next performed to determine the SD in %GIA. For this analysis, %GIA was used as a response variable, while mAb choice and test concentrations were used as explanatory variables. Total variance may be divided into two parts, the first being “signal” (i.e. the variance that can be explained by which mAb was tested at what concentration in the GIA), which a researcher actually wants to measure, and the second being EoA (i.e. the remaining variance). The proportions of signal and EoA were 89 % and 11 % respectively (**Figure 3A**). Based on the variance of EoA, the SD in %GIA of the assay was calculated as 13.1, which was close to the average SD of 12.2 determined earlier (**Figure 1A**), as predicted. In the previously mentioned publication investigating the EoA of *Pf*GIA [25], the SD was given as 7.5. Since the estimated SD value in *Pk*GIA with anti-*Pv*DBPII mAbs was ∼1.7 times higher, to investigate whether the two SD values were truly different, the 95 percent confidence interval (95%CI) of SD for the *Pk*GIA was estimated by an assay-stratified bootstrapping method. Resulting from this, the 95%CI of SD was 8.4 to 15.7, which suggests that the EoA in the *Pk*GIA might be slightly larger than the EoA in the *Pf*GIA, but not radically different. Utilizing the SD value of 13.1, the impact of repeat assay on the EoA in %GIA was investigated (**Figure 3B**). When a sample is tested in a single assay, the 95%CI of the %GIA is shown to be +/-25.7 % points of observed %GIA. If the 95%CI is to be narrowed down to +/-15 % points, three assays are required; whereas, when striving for a 95%CI of +/-10 % points, four additional assays (i.e., a total of seven assays) will be needed. After this, performing one more extra assay only further reduces the 95% CI width by < 2 % points.

**Figure 3:**
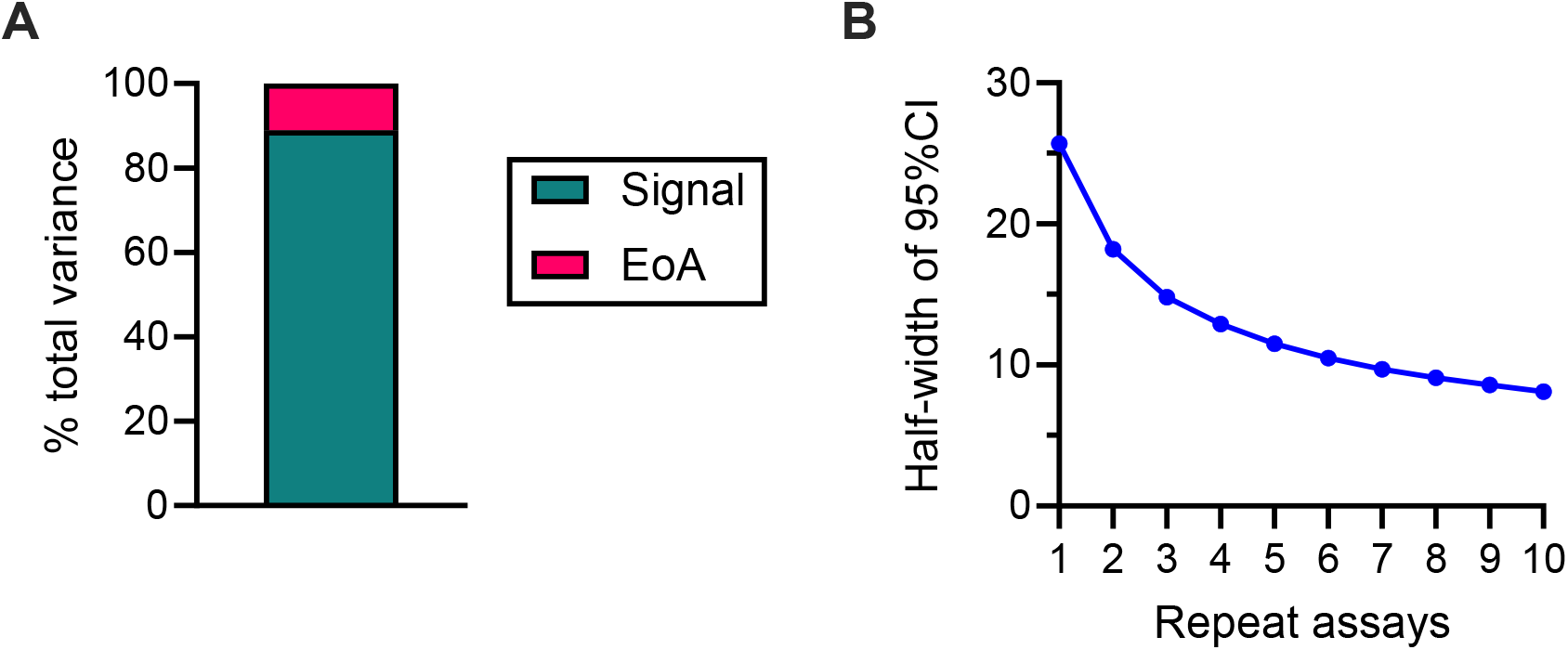
Range of error in %GIA estimates. (**A**) A linear regression analysis was performed with %GIA as a response variable, and mAb of choice as well as test concentrations as explanatory variables. A proportion (%) of variance explained by mAb and test concentration (Signal) and that for residual (EoA) in the total variance are presented. (**B**) From the standard deviation (SD) value of 13.1, the half width of 95%CI in %GIA was calculated to determine impact for a given number of repeat assays.

Subsequently, the SD for the antibody concentrations that gave 50, 40 or 30 percent of growth inhibition (GIA_50_/GIA_40_/GIA_30_, respectively) was determined. The mAb DB6 could not facilitate more than 50 %GIA in five out of ten assays, even at the maximum concentration of 2 mg/mL. Consequently, DB6 data were excluded from analysis in the GIA_50_ readout (while DB6 data were included for GIA_40_/GIA_30_ analysis). With only three (for GIA_50_ data) or four (GIA_40_ and GIA_30_) data points (one average, one SD and one %CV value per mAb) it was difficult to construct and evaluate a proper model as was done for **Figure 1**. Hence, the assumption was made that a constant SD model would reasonably be able to explain log-transformed GIA_50_, GIA_40_ and GIA_30_ values (LogGIA_50_, LogGIA_40_, and LogGIA_30_, respectively), as was shown for the *Pf*GIA in the aforementioned publication [25], where SD of LogGIA_50_ was relatively stable regardless of LogGIA_50_ level, while the SD of non-transformed GIA_50_ was affected by LogGIA_50_ level.

Making use of linear regression models once more, SDs in LogGIA_50_ (excluding DB6 data), LogGIA_40_ and LogGIA_30_ (both including DB6 data) were calculated and the 95%CI of the SDs again determined by a bootstrap analysis (**Figure 4A**). 95%CI ranges in SD exhibited an overlap in all three readouts, indicating that the EoA is similar when GIA_50_, GIA_40_ or GIA_30_ values are used for analysis. For the LogGIA_50_ readout, the SD was calculated as 0.299 and this value was used to investigate the impact of repeat assays on the EoA in a non-transformed GIA_50_ (**Figure 4B**). When testing a sample in a single assay (at serial dilutions), where the observed GIA_50_ is 1 mg/mL, the 95%CI is between 0.3 to 3.9 mg/mL. If three repeated assays are performed, where the geometric mean of GIA_50_ = 1 mg/mL, the 95%CI range narrows to 0.5 to 2.2 mg/mL, while with an additional seven assays (a total of ten assays) the 95%CI range becomes 0.7 to 1.5 mg/mL.

**Figure 4:**
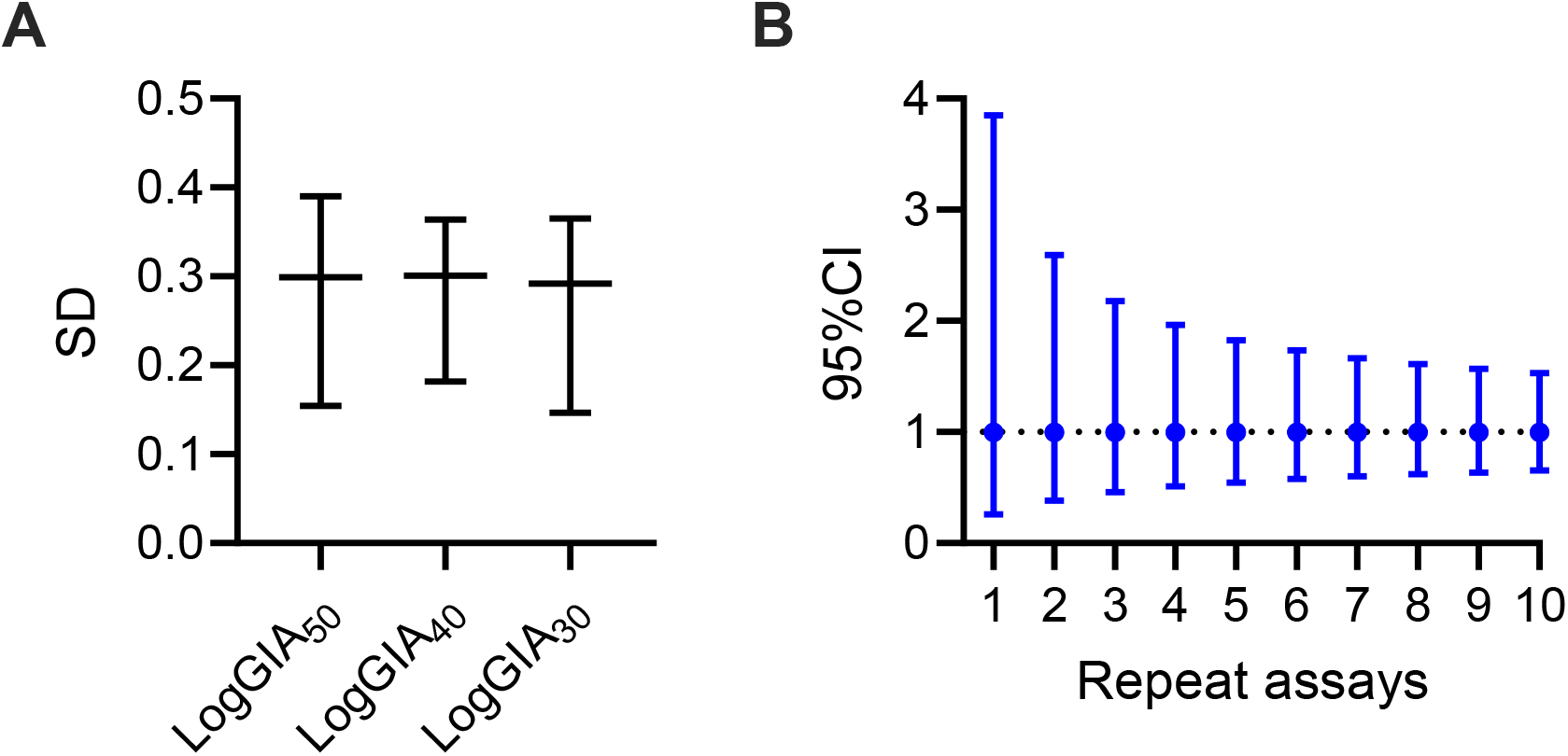
Assay variation in LogGIA_50_/GIA_40_/GIA_30_. (**A**) For each mAb in each assay, the concentrations that gave 50, 40 or 30 %GIA were calculated (GIA_50_/GIA_40_/GIA_30_, respectively). Subsequently, linear regression analyses were performed, with Log-transformed GIA_50_/GIA_40_/GIA_30_ values as a response variable and mAb tested as an explanatory variable. The SD in LogGIA_50_/GIA_40_/GIA_30_ was calculated from the residual variance. The 95%CI of SD for each readout was estimated from assay-stratified bootstrap analysis. The SD of LogGIA_50_ was calculated from three mAbs (excluding DB6), while those for LogGIA_40_ and LogGIA_30_ were calculated from all four mAbs. (**B**)The 95%CI range for a number of repeated assays is shown in a non-transformed GIA_50_ scale when the observed geometric mean of GIA_50_ = 1 mg/mL.

All of the data evaluated so far made use of monoclonal antibodies only. For investigating the EoA in *Pk*GIAs conducted with human polyclonal antibodies (pAbs), we turned to the analysis of some of our recently published data from a *Pv*DBPII Phase 1/2a clinical trial, involving CHMI. In this study, for which the *Pk*GIAs were conducted at the Laboratory of Malaria and Vector Research (LMVR), 80 human anti-*Pv*DBPII pAbs were tested at a single concentration of 10 mg/mL in three independent assays using three different batches of RBCs [19]. The original GIA values can be found in supplementary **Table S1**. Similar to what was found for the mAb dataset accrued in Oxford, for the pAb data, a constant SD model was more appropriate for subsequent analysis when compared to a constant %CV model (Spearman’s rank correlation coefficient *p* = 0.4246 vs *p* < 0.0001) (**Figure 5A, B**). Just like for the mAb data, a linear regression analysis was performed where %GIA value was utilized as a response variable and pAb as an explanatory variable. Based on the analysis, the SD in %GIA for *Pk*GIA conducted at LMVR using pAb was estimated as 5.94. Once more, the impact of repeat assays on the 95%CI was evaluated (**Figure 5C**). The 95%CI range shrinks from +/-11.6 % points for a single assay to +/-6.7 % points for three repeats, while after 10 assays are performed the range is estimated to be at +/-3.7 % points.

**Figure 5:**
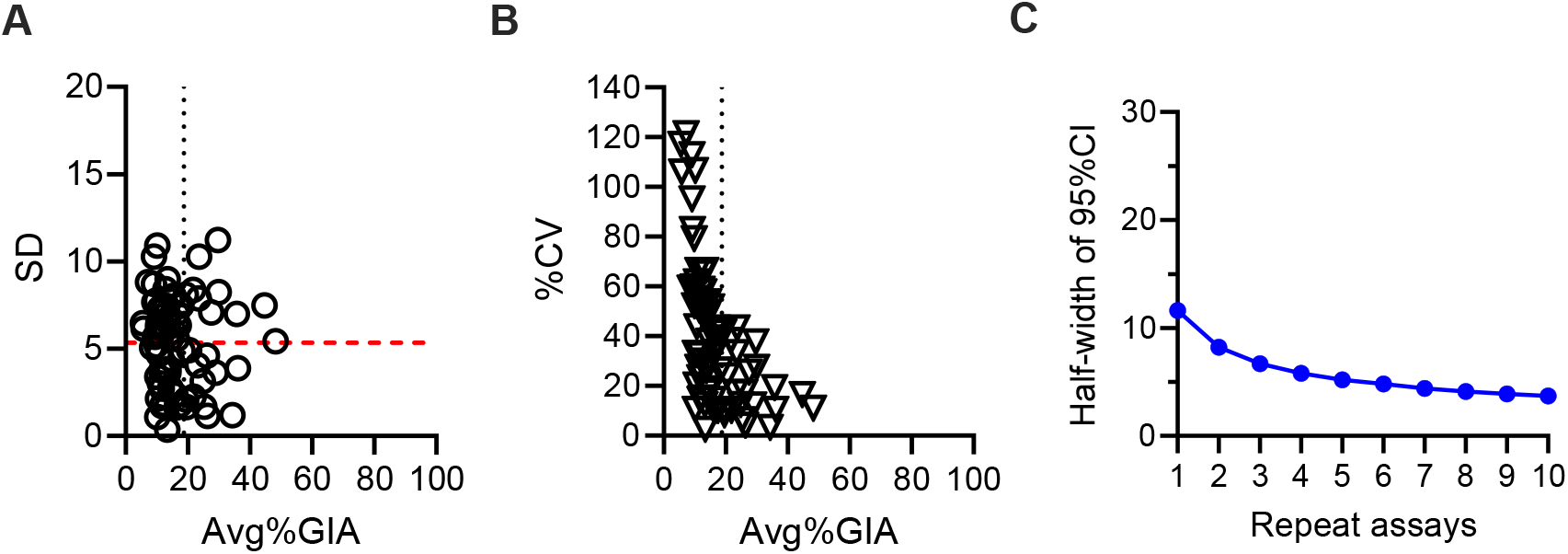
EoA in *Pk*GIA from a different dataset with human polyclonal antibodies conducted at LMVR. Eighty human anti-*Pv*DBPII polyclonal antibodies were evaluated at a concentration of 10 mg/mL in three independent GIAs using three different batches of RBCs. For each sample, Avg, SD (**A**) and %CV (**B**) were calculated. The vertical black line in panels **A** and **B** indicates the mean +2 SD value for 0.5 mg/mL EBL040 negative control mAb tested at the University of Oxford; the horizontal red line in **A** demonstrates the mean SD (5.34) of all data points. (**C**) The half-width of the 95%CI in %GIA for a given number of repeat assays is shown.

In the *Pk*GIA at the LMVR, relatively lower %GIA values were observed for the pAb compared to the mAb dataset tested at the University of Oxford. When results of the negative control EBL040 mAb tested at Oxford were used to determine a threshold of positive response (mean plus two SD, 18.8 %GIA), only 22 out of 80 samples (i.e. 27.5 %) exhibited positive Avg%GIA. To confirm that the SD value was stable regardless of positive or negative responses, another linear regression analysis was executed utilizing only the data from the 22 positive samples. The resulting SD value was determined to be 6.03, and thus very close to the SD of 5.94 from the analysis of the whole dataset.

## 4. Discussion

This is the first study that investigates EoA in GIA for transgenic *Pk* parasites expressing *Pv*DBP, instead of their native *Pk*DBPα (*Pv*DBP^OR^/Δβγ). Our *Pk*GIA data made use of four human anti-*Pv*DBPII mAbs tested at different concentrations, as well as eighty human vaccine-induced anti-*Pv*DBPII polyclonal antibodies at 10 mg/mL. In both cases, (non-transformed) %GIA data were explained better by a constant SD model than a constant %CV model. The SD in %GIA readout for the mAb dataset was 13.1 and 5.94 for the pAb dataset. In addition, based on the mAb data, the SD of LogGIA_50_ was calculated as 0.299. Using the SD values, the impact of repeat assays on the error range (95%CI) of observed %GIA or GIA_50_ values were estimated.

Similar to the previous study, where the EoA was evaluated in *Pf*GIA using anti-RH5 human antibodies [25], we also observed a significant assay-to-assay variation in *Pk*GIA with anti-*Pv*DBPII mAbs in this study. This finding emphasizes the difficulty with directly comparing GIA results from different investigations, especially when the results from only a single or two repeat assays are reported. The 95%CI ranges calculated in this study for a given number of assays will help not only in comparing different formulations and/or immunological strategies to develop the second generation of *Pv*DBP-based vaccines, but will also provide researchers with insight on how to interpret GIA results from different studies.

In our analysis, there was an almost two-fold difference in the best estimated SD in %GIA for *Pk*GIA conducted in the two examined laboratories (13.1 at the University of Oxford vs 5.94 at the LMVR). Hence, one might wonder whether researchers and vaccine developers need to use different SD values and 95%CI ranges (which in turn are calculated from the SD values), depending on the sample type (either mAb or pAb) or laboratory where the *Pk*GIA is performed. Precision and accuracy of an assay may vary, particularly between laboratories [28]. With the ever increasing need for international collaboration between laboratories for facilitation of development of effective malaria vaccines [29], this interlaboratory assay variability is a considerable factor when it comes to interpretation of results, as underlined by our study. Interlaboratory variability may arise from different materials, reagents and kits used, small differences in culture (e.g. gas mixture, incubation times) and environmental conditions (e.g. collection and storage procedures, room or storage temperature or light conditions with a photometric assay). Moreover, assay protocols may vary and even if an identical protocol is used, divergent routines within the boundaries of the protocol might have come about. Another aspect to keep in mind are operator handling and training, including technical proficiency and day-to-day performance [30] as well as detection and analysis methods [28]. Be that as it may, our assay-stratified bootstrap analysis also raised the question as to whether the two SD values are truly different. Based on this analysis, the 95%CI for Oxford’s SD was between 8.4 to 15.7, i.e. the true SD value of the Oxford %GIA data falls with 95% probability – and thus highly likely – anywhere between 8.4 to 15.7. Likewise, the observed SD value for the LMVR data (5.94) could naturally deviate from the true SD value to a certain degree. At the University of Oxford, ten assays were conducted, thus we are of the opinion that performing an assay-stratified bootstrap analysis from 1010 possible data combinations is reasonable to estimate the 95%CI range of the true SD value. On the other hand, only three assays per sample were performed at the LMVR, therefore the same analysis was not performed (as there are only 33 possible combinations), and we could not assess whether the two SD values were of significant difference by a statistical test. To answer whether the SD values from the two data sets are truly divergent or not, further investigation is required, ideally with both laboratories performing additional *Pk*GIAs with the same variety of samples in multiple assays. Interestingly, the previously reported SD in %GIA for *Pf*GIA (at the LMVR) was between the two SD values reported in this study (SD = 7.5) [25], although the *Pf*GIA used a different species of parasite for GIA to test antibodies against a different antigen. Therefore, whilst it is possible that EoA in GIA could be dependent on the parasites employed (e.g. different *Pf* strains or transgenic *Pk* parasites), target antigen(s) and/or laboratory, unless experimentally confirmed, it might be acceptable to assume that the SD in %GIA is around 10, at least when the GIA is performed with reasonably strict adherence to the same protocols and procedures, as in the two investigated laboratories here.

The evaluated mAb data included blood from donors with different Duffy blood group (or DARC/Fy) serophenotypes. The two codominant DARC alleles *FY*A* and *FY*B* encode the Fy^a^ or Fy^b^ antigens resulting in the Fy^a+/b+^, Fy^a-/b+^ and Fya+/b-phenotypes being expressed on RBCs [31]. Thus, four main Duffy serophenotypes exist of varying frequency depending on geolocation [32] and ethnicity (within a mixed-descent population in the same location) [33]. As mentioned before, the DARC genotype plays a significant role in *Pv* RBC invasion, with large populations in Sub-Saharan (and particularly West) Africa resistant to *Pv* infection due to being of Fy^a-/b-^(or “Duffy negative”) serophenotype [7,32]. Naturally, this would seem to reciprocally correlate with a low incidence of *Pv* malaria and might have prevented endemicity of *Pv* in this region of the world [34]. Yet recently conflicting evidence has come to light [35,36], reporting higher *Pv* malaria incidence in Africa than originally thought [37], especially in regions where *Pf* burden has been lowered [4]. Due to the emergence and spreading of *Pv* strains with the apparent ability to invade Duffy negative RBCs [38], it has been proposed that there could be a hidden transmission in Fy^a-/b-^populations in Africa. These cases generally show lower parasitaemia and may thus contribute to the undetected low-transmission reservoir in Africa [39]. Very recently, two investigations could show the existence of a subset of Fy^a-/b-^erythroblasts that transiently express DARC and can be infected by *Pv*, thus providing a scientific reasoning behind the transmission of *Pv* infection in Duffy negative individuals [40,41]. With the *Pv* invasion pathway so reliant on the specific DARC genotype, it could be speculated that this may also have a significant effect on *Pk*GIA results. However, in our study, only Duffy-positive RBCs were utilized, as the transgenic *Pv*DBPOR/Δβγ parasite is known to only infect Duffy-positive, but not Duffy-negative, RBCs (as expected) [23]. Our results did not demonstrate an obvious Fy effect on ΔAve%GIA above the assay-to-assay variation within the same Fy serophenotype. In other words, anti-*Pv*DBPII mAbs used in this study were equally inhibitory and the same SD (or 95%CI range) could be used to interpret the *Pk*GIA results for all three Fy positive (i.e. Fy^a+/b+^, Fy^a+/b-^, Fy^a-/b+^) serophenotypes. In summary, Fy serophenotype (at least if it is positive) may have a minimum or no impact on %GIA measured by the *Pk*GIA, while the impact of Fy serophenotype on anti-*Pv*DBP vaccine efficacy needs to be more fully evaluated in future larger Phase 2 trials.

In the aforementioned *Pf*GIA study, which exhibited a “balanced” design (i.e., the same set of samples were tested with multiple RBCs on each day, and the assays were repeated on multiple days), it was possible to separate out RBC-to-RBC variation (on the same day) and day-to-day variation (within the same RBC batch). It was shown that the RBC donor effect was approximately four times higher than the day effect on EoA. However, in this study only one RBC batch was used on one assay day for most of the data sets analyzed. Hence, while the linear regression analysis for the Oxford data did show a significant assay-to-assay variation (*p* < 0.001) in ΔAve%GIA, we could not evaluate how much assay-to-assay variation could be explained by RBC-to-RBC or day-to-day variation. Interestingly, previous studies have found variations in *Pk* growth rate when blood samples drawn from different donors were used for *in vitro* culture. These appear to be largely independent of DARC phenotype, suggesting that blood phenotypes beyond DARC and even donor-specific factors (e.g. diet, health, medication) potentially impact this variability in growth rate [21,23], which may in turn have an effect on GIA readouts. Another source for variability in the assay might be the state of the cultured parasites. A key variable for many examinations involving *Plasmodium* is the specific condition the parasites are in on the particular day of the experiment and the days preceding it. To investigate this in detail, further work with specifically designed experiments would be required. Nonetheless, the inclusion of a “parasite health criterion” or establishing the protocolization of parasite preparatory stages might aid in further reducing the SD both in the GIA and other similar assays involving *Plasmodium*.

In GIA studies, not only %GIA of test samples at the same concentration(s), but also GIA_50_ values, have been widely used to compare functional activity among different samples. However, the determination of the 50 %GIA threshold, instead of 60 or 30 %GIA for example, is chosen rather arbitrarily. In addition, although a significant correlation between *in vivo* growth inhibition (IVGI) and *in vitro* growth inhibition (%GIA) was observed in the previously mentioned *Pv*DBPII Phase 1/2a vaccine trials [19], the level of %GIA was generally low at 10 mg/mL (**Figure 5A**). Therefore, using mAb *Pk*GIA data, we explored the possibility of other readouts, namely of GIA_40_ and GIA_30_, for future studies. As shown in **Figure 4A**, SDs for all three readouts were similar, indicating that the GIA_40_ or GIA_30_ readout could be used to compare different samples with similar precision as the GIA_50_ readout. Of note, the 95%CI of SD for LogGIA_50_ from this study was between 0.154 and 0.390, which overlapped with the best estimate of SD for LogGIA_50_ in *Pf*GIA reported before (0.206) [25]. The result again suggests that the EoA in the *Pk*GIA conducted at the University of Oxford and the EoA in the *Pf*GIA conducted at the LMVR might be of similar magnitude.

Performing multiple assays naturally improves the reliance of the accrued results. Under the assumption that a constant SD model reasonably explains the *Pk*GIA results, the shrinkage of the 95%CI with repeat assays in our experiments was not linear, i.e. the 95%CI window shrinks more from assay 1 to assay 2 than from assay 2 to 3 and so forth, while there is almost no diminishment from assay 9 to 10, as seen in **Figures 3B, 4B** and **5C**. Depending on the assay precision required, while also keeping practicality in mind, the number of repeat assays should be optimized in each study. For our data, it might not be worthwhile to perform more than four to five repeat assays for the purpose of minimizing the 95%CI window.

There are several limitations to our study. Only one species of parasite and strain was employed in the assay, namely A1-H.1 *Pk* transgenic for Sal-1 *Pv*DBP, to evaluate antibodies against only one target molecule region, *Pv*DBPII. Furthermore, relative parasitaemia was determined by pLDH activity in both laboratories. Parasite species or strain, target antigen, and/or a method of parasitaemia determination in the GIA – of which a multitude exists, e.g. biochemical assays like the pLDH assay, or assays based on microscopy or flow cytometry [42] – may influence the EoA and thus, upcoming evaluations should investigate on these parameters. With our *Pk*GIA and previous *Pf*GIA studies taken together, a SD of 10 in %GIA and SD of 0.2 - 0.3 in LogGIA_50_ can be considered reasonable starting points to design an EoA determination experiment for different GIAs in the future. Moreover, the source of the EoA was not assessed in our experiments here. In addition to the RBC-to-RBC and day-to-day variations discussed above, well-to-well, plate-to-plate and operator-to-operator variations could contribute to the final assay-to-assay variation. Well-to-well variation has been calculated for the Oxford dataset, it was relatively small compared to the entire assay-to-assay variation (Median SD = 3.95, see **Supplemental Figure S1**); yet, we do not possess enough data to estimate the other sources of variation mentioned above. A future “balanced” study with a higher number of assays (using multiple batches of RBCs) and multiple samples will help in identifying the source(s) of the *Pk*GIA EoA.

## 5. Conclusions

With recent data underlining the increasing importance of *P. vivax* control globally, the calls for an efficacious vaccine against this parasite species will only become more urgent. Robust candidate selection tools will be required for achieving development of such a vaccine. The GIA with transgenic *Pk* parasites expressing target *Pv* antigens has great potential in filling this role for blood-stage candidate vaccines, particularly with *Pk*GIA results correlating with *in vivo* protection post-CHMI. This investigation marks the first study investigating EoA in the *Pk*GIA, in which significant assay-to-assay variation was observed. These results might be considered in the down-selection process of new candidate formulations and thus aid in the development of a novel blood-stage vaccine, especially one with *Pv*DBP as its target.

## Supplementary Materials

Figure S1: Intra-assay variability in %GIA in the mAb dataset accrued at the University of Oxford; Table S1: Original %GIA data for each sample at each concentration in every assay at both the University of Oxford and the LMVR.

## Author Contributions

“Conceptualization, J.E.M., S.J.D., and K.M.; methodology, J.E.M., S.J.D., and K.M.; software, J.E.M. and K.M.; validation, J.E.M., S.J.D., and K.M.; formal analysis, J.E.M. and K.M.; investigation, J.E.M., C.A.R., M.B., D.Q., M.M.H., A.D., C.A.L. and K.M.; resources, S.E.S., C.E.C, A.M.M, and R.W.M; data curation, J.E.M. and K.M.; writing—original draft preparation, J.E.M. and K.M.; writing—review and editing, J.E.M., S.J.D. and K.M..; visualization, J.E.M. and K.M.; supervision, S.J.D. and K.M. All authors have read and agreed to the published version of the manuscript.”

## Funding

During his overseas stay in the United Kingdom, J.E.M. was supported by research scholarships from the German Academic Exchange Service (DAAD) and the University of Hamburg. A.D., C.A.L. and K. M. were supported by the intramural program of the NIAID/NIH. S.J.D. is a Jenner Investigator and held a Wellcome Trust Senior Fellowship (106917/Z/15/Z). R.W.M is supported by a Wellcome Trust Discovery Award (225844/Z/22/Z) as well as the OptiViVax consortium. This work was furthermore supported in part by OptiViVax; this project is co-funded from the European Union Horizon Europe programme under grant agreement No. 101080744 and the project also receives funding from UK Research and Innovation (UKRI) under the UK government’s Horizon Europe funding guarantee (grant No. 10077974). This work was also supported in part by the National Institute for Health Research (NIHR) Oxford Biomedical Research Centre (BRC) and NHS Blood & Transplant (NHSBT; who provided material), the views expressed are those of the authors and not necessarily those of the NIHR or the Department of Health and Social Care or NHSBT.

## Institutional Review Board Statement/Informed Consent Statement

All of the blood donations for laboratory assays at the University of Oxford are covered under ethical approval from the United Kingdom’s National Health Service Research Ethics Services (Ref 18/LO/0415, protocol number OVC002). The clinical trials received ethical approval from UK National Health Service Research Ethics Services, (VAC069: Hampshire A Research Ethics Committee, Ref 18/SC/0577; VAC071: Oxford A Research Ethics Committee, Ref 19/SC/0193; VAC079: Oxford A Research Ethics Committee, Ref 19/SC/0330). The vaccine trials were also approved by the UK Medicines and Healthcare products Regulatory Agency (VAC071: EudraCT 2019-000643-27; VAC079: EudraCT 2019-002872-14). All participants provided written informed consent and the trials were conducted according to the principles of the current revision of the Declaration of Helsinki 2008 and ICH guidelines for Good Clinical Practice.

## Data Availability Statement

The data presented in this study are available in Supplementary Table S1. Other parts of data which support the findings of the clinical trial(s) and/or specific reagents are available on request from the corresponding authors.

## Acknowledgments

The authors would like to express their thanks to the various blood donors and participants in the clinical trials. We also thank Tom Rawlinson, Kirsty McHugh, Jenny Bryant, Kate Skinner, Amy Boyd, Patries Fisher and Lana Strmecki (University of Oxford) for assistance and appreciate Michael Fay (NIAID/NIH) for statistical advice.

## Conflicts of Interest

C.E.C. is an inventor on patents that relate to binding domains of erythrocytebinding proteins of *Plasmodium* parasites including *Pv*DBP (patent no. 6962987; binding domains from *Plasmodium vivax* and *Plasmodium falciparum* erythrocyte binding proteins).

## Notes

### Competing Interest Statement

C.E.C. is an inventor on patents that relate to binding domains of erythrocyte-binding proteins of *Plasmodium* parasites including *Pv*DBP (patent no. 6962987; binding domains from *Plasmodium vivax* and *Plasmodium falciparum* erythrocyte binding proteins).

## References

1. World Malaria Report 2023; World Health Organization: Geneva, 2023; ISBN 978-92-4-008617-3.

2. Gething, P.W.; Elyazar, I.R.F.; Moyes, C.L.; Smith, D.L.; Battle, K.E.; Guerra, C.A.; Patil, A.P.; Tatem, A.J.; Howes, R.E.; Myers, M.F.; et al. A Long Neglected World Malaria Map: Plasmodium Vivax Endemicity in 2010. PLoS Negl Trop Dis 2012, 6, e1814, doi:10.1371/journal.pntd.0001814.

3. Mueller, I.; Galinski, M.R.; Baird, J.K.; Carlton, J.M.; Kochar, D.K.; Alonso, P.L.; del Portillo, H.A. Key Gaps in the Knowledge of Plasmodium Vivax, a Neglected Human Malaria Parasite. Lancet Infect Dis 2009, 9, 555–566, doi:10.1016/S1473-3099(09)70177-X.

4. Price, R.N.; Commons, R.J.; Battle, K.E.; Thriemer, K.; Mendis, K. Plasmodium Vivax in the Era of the Shrinking P. Falciparum Map. Trends Parasitol 2020, 36, 560–570, doi:10.1016/j.pt.2020.03.009.

5. White, N.J. Determinants of Relapse Periodicity in Plasmodium Vivax Malaria. Malaria Journal 2011, 10, 297, doi:10.1186/1475-2875-10-297.

6. White, M.; Amino, R.; Mueller, I. Theoretical Implications of a Pre-Erythrocytic Plasmodium Vivax Vaccine for Preventing Relapses. Trends Parasitol 2017, 33, 260–263, doi:10.1016/j.pt.2016.12.011.

7. Miller, L.H.; Mason, S.J.; Clyde, D.F.; McGinniss, M.H. The Resistance Factor to Plasmodium Vivax in Blacks. The Duffy-Blood-Group Genotype, FyFy. N Engl J Med 1976, 295, 302–304, doi:10.1056/NEJM197608052950602.

8. Galinski, M.R.; Medina, C.C.; Ingravallo, P.; Barnwell, J.W. A Reticulocyte-Binding Protein Complex of Plasmodium Vivax Merozoites. Cell 1992, 69, 1213–1226, doi:10.1016/0092-8674(92)90642-p.

9. Battle, K.E.; Lucas, T.C.D.; Nguyen, M.; Howes, R.E.; Nandi, A.K.; Twohig, K.A.; Pfeffer, D.A.; Cameron, E.; Rao, P.C.; Casey, D.; et al. Mapping the Global Endemicity and Clinical Burden of Plasmodium Vivax, 2000–17: A Spatial and Temporal Modelling Study. The Lancet 2019, 394, 332–343, doi:10.1016/S0140-6736(19)31096-7.

10. Baird, J.K. Evidence and Implications of Mortality Associated with Acute Plasmodium Vivax Malaria. Clin Microbiol Rev 2013, 26, 36–57, doi:10.1128/CMR.00074-12.

11. Galinski, M.R.; Barnwell, J.W. Plasmodium Vivax: Who Cares? Malar J 2008, 7 Suppl 1, S9, doi:10.1186/1475-2875-7-S1-S9.

12. Draper, S.J.; Sack, B.K.; King, C.R.; Nielsen, C.M.; Rayner, J.C.; Higgins, M.K.; Long, C.A.; Seder, R.A. Malaria Vaccines: Recent Advances and New Horizons. Cell Host Microbe 2018, 24, 43–56, doi:10.1016/j.chom.2018.06.008.

13. McCarthy, J.S.; Griffin, P.M.; Sekuloski, S.; Bright, A.T.; Rockett, R.; Looke, D.; Elliott, S.; Whiley, D.; Sloots, T.; Winzeler, E.A.; et al. Experimentally Induced Blood-Stage Plasmodium Vivax Infection in Healthy Volunteers. J Infect Dis 2013, 208, 1688–1694, doi:10.1093/infdis/jit394.

14. Minassian, A.M.; Themistocleous, Y.; Silk, S.E.; Barrett, J.R.; Kemp, A.; Quinkert, D.; Nielsen, C.M.; Edwards, N.J.; Rawlinson, T.A.; Ramos Lopez, F.; et al. Controlled Human Malaria Infection with a Clone of Plasmodium Vivax with High-Quality Genome Assembly. JCI Insight 2021, 6, e152465, doi:10.1172/jci.insight.152465.

15. Payne, R.O.; Silk, S.E.; Elias, S.C.; Milne, K.H.; Rawlinson, T.A.; Llewellyn, D.; Shakri, A.R.; Jin, J.; Labbé, G.M.; Edwards, N.J.; et al. Human Vaccination against Plasmodium Vivax Duffy-Binding Protein Induces Strain-Transcending Antibodies. JCI Insight 2017, 2, e93683, 93683, doi:10.1172/jci.insight.93683.

16. Singh, K.; Mukherjee, P.; Shakri, A.R.; Singh, A.; Pandey, G.; Bakshi, M.; Uppal, G.; Jena, R.; Rawat, A.; Kumar, P.; et al. Malaria Vaccine Candidate Based on Duffy-Binding Protein Elicits Strain Transcending Functional Antibodies in a Phase I Trial. npj Vaccines 2018, 3, 1–10, doi:10.1038/s41541-018-0083-3.

17. Batchelor, J.D.; Zahm, J.A.; Tolia, N.H. Dimerization of Plasmodium Vivax DBP Is Induced upon Receptor Binding and Drives Recognition of DARC. Nat Struct Mol Biol 2011, 18, 908–914, doi:10.1038/nsmb.2088.

18. Batchelor, J.D.; Malpede, B.M.; Omattage, N.S.; DeKoster, G.T.; Henzler-Wildman, K.A.; Tolia, N.H. Red Blood Cell Invasion by Plasmodium Vivax: Structural Basis for DBP Engagement of DARC. PLoS Pathog 2014, 10, e1003869, doi:10.1371/journal.ppat.1003869.

19. Hou, M.M.; Barrett, J.R.; Themistocleous, Y.; Rawlinson, T.A.; Diouf, A.; Martinez, F.J.; Nielsen, C.M.; Lias, A.M.; King, L.D.W.; Edwards, N.J.; et al. Vaccination with Plasmodium Vivax Duffy-Binding Protein Inhibits Parasite Growth during Controlled Human Malaria Infection. Sci Transl Med 2023, 15, eadf1782, doi:10.1126/scitranslmed.adf1782.

20. Rawlinson, T.A.; Barber, N.M.; Mohring, F.; Cho, J.S.; Kosaisavee, V.; Gérard, S.F.; Alanine, D.G.W.; Labbé, G.M.; Elias, S.C.; Silk, S.E.; et al. Structural Basis for Inhibition of Plasmodium Vivax Invasion by a Broadly Neutralizing Vaccine-Induced Human Antibody. Nat Microbiol 2019, 4, 1497–1507, doi:10.1038/s41564-019-0462-1.

21. Mohring, F.; Hart, M.N.; Rawlinson, T.A.; Henrici, R.; Charleston, J.A.; Diez Benavente, E.; Patel, A.; Hall, J.; Almond, N.; Campino, S.; et al. Rapid and Iterative Genome Editing in the Malaria Parasite Plasmodium Knowlesi Provides New Tools for P. Vivax Research. Elife 2019, 8, e45829, doi:10.7554/eLife.45829.

22. Mohring, F.; Rawlinson, T.A.; Draper, S.J.; Moon, R.W. Multiplication and Growth Inhibition Activity Assays for the Zoonotic Malaria Parasite, Plasmodium Knowlesi. Bio Protoc 2020, 10, e3743, doi:10.21769/BioProtoc.3743.

23. Moon, R.W.; Hall, J.; Rangkuti, F.; Ho, Y.S.; Almond, N.; Mitchell, G.H.; Pain, A.; Holder, A.A.; Blackman, M.J. Adaptation of the Genetically Tractable Malaria Pathogen Plasmodium Knowlesi to Continuous Culture in Human Erythrocytes. Proc Natl Acad Sci U S A 2013, 110, 531–536, doi:10.1073/pnas.1216457110.

24. Watson, Q.D.; Carias, L.L.; Malachin, A.; Redinger, K.R.; Bosch, J.; Bardelli, M.; Baldor, L.; Feufack-Donfack, L.B.; Popovici, J.; Moon, R.W.; et al. Human Monoclonal Antibodies Inhibit Invasion of Transgenic Plasmodium Knowlesi Expressing Plasmodium Vivax Duffy Binding Protein. Malar J 2023, 22, 369, doi:10.1186/s12936-023-04766-1.

25. Miura, K.; Diouf, A.; Fay, M.P.; Barrett, J.R.; Payne, R.O.; Olotu, A.I.; Minassian, A.M.; Silk, S.E.; Draper, S.J.; Long, C. A. Assessment of Precision in Growth Inhibition Assay (GIA) Using Human Anti-PfRH5 Antibodies. Malar J 2023, 22, 159, doi:10.1186/s12936-023-04591-6.

26. Miura, K.; Zhou, H.; Diouf, A.; Moretz, S.E.; Fay, M.P.; Miller, L.H.; Martin, L.B.; Pierce, M.A.; Ellis, R.D.; Mullen, G.E.D.; et al. Anti-Apical-Membrane-Antigen-1 Antibody Is More Effective than Anti-42-Kilodalton-Merozoite-Surface-Protein-1 Antibody in Inhibiting Plasmodium Falciparum Growth, as Determined by the in Vitro Growth Inhibition Assay. Clin Vaccine Immunol 2009, 16, 963–968, doi:10.1128/CVI.00042-09.

27. Rijal, P.; Elias, S.C.; Machado, S.R.; Xiao, J.; Schimanski, L.; O’Dowd, V.; Baker, T.; Barry, E.; Mendelsohn, S.C.; Cherry, C.J.; et al. Therapeutic Monoclonal Antibodies for Ebola Virus Infection Derived from Vaccinated Humans. Cell Rep 2019, 27, 172–186.e7, doi:10.1016/j.celrep.2019.03.020.

28. Knight, V.; Long, T.; Meng, Q.H.; Linden, M.A.; Rhoads, D.D. Variability in the Laboratory Measurement of Cytokines: A Longitudinal Summary of a College of American Pathologists Proficiency Testing Survey. Arch Pathol Lab Med 2020, 144, 1230–1233, doi:10.5858/arpa.2019-0519-CP.

29. Birkett, A.J.; Moorthy, V.S.; Loucq, C.; Chitnis, C.E.; Kaslow, D.C. Malaria Vaccine R&D in the Decade of Vaccines: Breakthroughs, Challenges and Opportunities. Vaccine 2013, 31 Suppl 2, B233–243, doi:10.1016/j.vaccine.2013.02.040.

30. Aziz, N.; Nishanian, P.; Fahey, J.L. Levels of Cytokines and Immune Activation Markers in Plasma in Human Immunodeficiency Virus Infection: Quality Control Procedures. Clin Diagn Lab Immunol 1998, 5, 755–761.

31. Dean, L. The Duffy Blood Group. In Blood Groups and Red Cell Antigens [Internet]; National Center for Biotechnology Information (US), 2005.

32. Howes, R.E.; Patil, A.P.; Piel, F.B.; Nyangiri, O.A.; Kabaria, C.W.; Gething, P.W.; Zimmerman, P.A.; Barnadas, C.; Beall, C.M.; Gebremedhin, A.; et al. The Global Distribution of the Duffy Blood Group. Nat Commun 2011, 2, 266, doi:10.1038/ncomms1265.

33. Nickel, R.G.; Willadsen, S.A.; Freidhoff, L.R.; Huang, S.-K.; Caraballo, L.; Naidu, R.P.; Levett, P.; Blumenthal, M.; Banks-Schlegel, S.; Bleecker, E.; et al. Determination of Duffy Genotypes in Three Populations of African Descent Using PCR and Sequence-Specific Oligonucleotides. Hum Immunol 1999, 60, 738–742, doi:10.1016/S0198-8859(99)00039-7.

34. Carter, R. Speculations on the Origins of Plasmodium Vivax Malaria. Trends Parasitol 2003, 19, 214–219, doi:10.1016/s1471-4922(03)00070-9.

35. Escalante, A.A.; Cornejo, O.E.; Freeland, D.E.; Poe, A.C.; Durrego, E.; Collins, W.E.; Lal, A.A. A Monkey’s Tale: The Origin of Plasmodium Vivax as a Human Malaria Parasite. Proc Natl Acad Sci U S A 2005, 102, 1980–1985, doi:10.1073/pnas.0409652102.

36. Howes, R.E.; Reiner, R.C.; Battle, K.E.; Longbottom, J.; Mappin, B.; Ordanovich, D.; Tatem, A.J.; Drakeley, C.; Gething, P.W.; Zimmerman, P.A.; et al. Plasmodium Vivax Transmission in Africa. PLoS Negl Trop Dis 2015, 9, e0004222, doi:10.1371/journal.pntd.0004222.

37. Twohig, K.A.; Pfeffer, D.A.; Baird, J.K.; Price, R.N.; Zimmerman, P.A.; Hay, S.I.; Gething, P.W.; Battle, K.E.; Howes, R.E. Growing Evidence of Plasmodium Vivax across Malaria-Endemic Africa. PLoS Negl Trop Dis 2019, 13, e0007140, doi:10.1371/journal.pntd.0007140.

38. Golassa, L.; Amenga-Etego, L.; Lo, E.; Amambua-Ngwa, A. The Biology of Unconventional Invasion of Duffy-Negative Reticulocytes by Plasmodium Vivax and Its Implication in Malaria Epidemiology and Public Health. Malar J 2020, 19, 299, doi:10.1186/s12936-020-03372-9.

39. Abate, A.; Bouyssou, I.; Mabilotte, S.; Doderer-Lang, C.; Dembele, L.; Menard, D.; Golassa, L. Vivax Malaria in Duffy-Negative Patients Shows Invariably Low Asexual Parasitaemia: Implication towards Malaria Control in Ethiopia. Malar J 2022, 21, 230, doi:10.1186/s12936-022-04250-2.

40. Bouyssou, I.; El Hoss, S.; Doderer-Lang, C.; Schoenhals, M.; Rasoloharimanana, L.T.; Vigan-Womas, I.; Ratsimbasoa, A.; Abate, A.; Golassa, L.; Mabilotte, S.; et al. Unveiling P. Vivax Invasion Pathways in Duffy-Negative Individuals. Cell Host Microbe 2023, S1931-3128(23)00457-2, doi:10.1016/j.chom.2023.11.007.

41. Dechavanne, C.; Dechavanne, S.; Bosch, J.; Metral, S.; Redinger, K.R.; Watson, Q.D.; Ratsimbasoa, A.C.; Roeper, B.; Krishnan, S.; Fong, R.; et al. Duffy Antigen Is Expressed during Erythropoiesis in Duffy-Negative Individuals. Cell Host Microbe 2023, S1931-3128(23)00450-X, doi:10.1016/j.chom.2023.10.019.

42. Duncan, C.J.A.; Hill, A.V.S.; Ellis, R.D. Can Growth Inhibition Assays (GIA) Predict Blood-Stage Malaria Vaccine Efficacy? Hum Vaccin Immunother 2012, 8, 706–714, doi:10.4161/hv.19712.

