## Supplementary Figure S1 for "Evaluation of Precision of the *Plasmodium knowlesi* Growth Inhibition Assay for *Plasmodium vivax* Duffy-Binding Protein-based Malaria Vaccine Development"

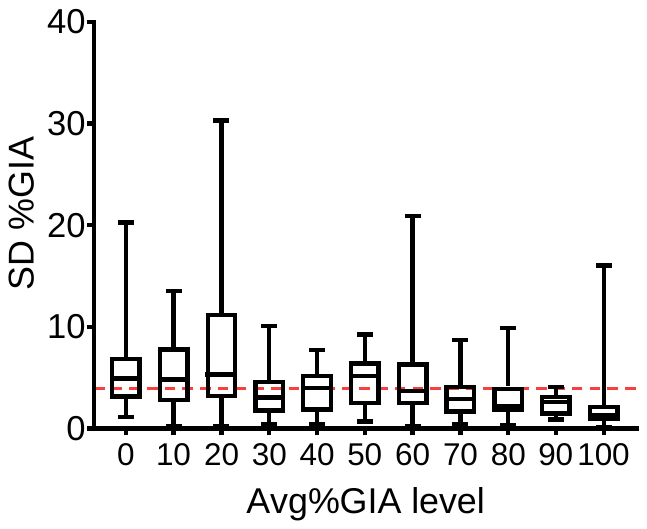

**Supplementary Figure S1: Intra-assay variability in %GIA in the mAb dataset accrued at the University of Oxford.** Average (Avg) and standard deviation (SD) of %GIA in duplicate or triplicate wells were calculated for each sample at each concentration in each assay (n = 2719). Thereafter, SD %GIA data were categorized by Avg%GIA levels in the following manner: SD %GIA data for each sample at each concentration were sorted into one of 11 bins (“0” to “100” for every tenth percentage point) based on the Avg%GIA level. The Avg%GIA level of “0” thus consists of all of the data from a sample or concentration with Avg%GIA values of < 5, while the level of “10” encompasses values from ≥ 5 to < 15, “20” contains values of ≥ 15 and < 25, and so forth. The Avg%GIA level of “100” encompasses all data with an Avg%GIA value of ≥ 95. For each Avg%GIA level, the box plot (of 25^th^, 50^th^ and 75^th^ percentiles) and error bars demonstrating the 2.5^th^ and 97.5^th^ percentile are shown. The horizontal red line indicates the average SD value of all data (SD = 3.95). This figure includes *Pk*GIA data not utilized for the main analysis shown in this article.
